# Using site-directed mutagenesis to further the understanding of insulin receptor-insulin like growth factor-1 receptor heterodimer structure

**DOI:** 10.1101/2024.03.05.583490

**Authors:** Samuel Turvey, Stephen P Muench, Tarik Issad, Colin WG Fishwick, Mark T Kearney, Katie J Simmons

## Abstract

Type 2 diabetes is characterised by the disruption of insulin and insulin-like growth factor (IGF) signalling. The key hubs of these signalling cascades - the Insulin receptor (IR) and Insulin-like growth factor 1 receptor (IGF1R) – are known to form functional IR-IGF1R hybrid receptors which are insulin resistant. However, the mechanisms underpinning IR-IGF1R hybrid formation are not fully understood, hindering the ability to modulate this for future therapies targeting this receptor. To pinpoint suitable sites for intervention, computational hotspot prediction was utilised to identify promising epitopes for targeting with point mutagenesis. Specific IGF1R point mutations F450A, R391A and D555A show reduced affinity of the hybrid receptor in a BRET based donor-saturation assay, confirming hybrid formation could be modulated at this interface. These data provide the basis for rational design of more effective hybrid receptor modulators, supporting the prospect of identifying a small molecule that specifically interacts with this target.

## Introduction

Over the past five decades changes in human lifestyle have contributed to an explosion of obesity^1^ and its frequent sequelae insulin resistant type 2 diabetes mellitus.^2^ A poorly understood hallmark of obesity and type 2 diabetes mellitus is disruption of insulin signalling,^3, 4^ leading to dysregulation of cellular growth and nutrient handling.^5^ The insulin receptor (IR) acts as a conduit for insulin-encoded information, which is transferred via a complex intracellular signalling network including the critical signalling nodes phosphatidylinositol 3-kinase (PI3-K) and the serine/threonine kinase Akt, to regulate cell metabolism.^6^ During evolution the IR and insulin-like growth factor-1 receptor (IGF1R) diverged from a single receptor in invertebrates,^7, 8^ into a more complex system in mammals.^9^ Stimulation of IR or IGF1R initiates phosphorylation of IR substrate (IRS) proteins at multiple tyrosine residues,^6^ phosphorylated IRS1 binds PI3-K initiating the conversion of the plasma lipid phosphatidylinositol 3,4,- bisphosphate to phosphatidylinositol 3,4,5-trisphosphate (PIP3) which activates the multifunctional serine−threonine kinase Akt.^10^ In endothelial cells Akt activates the endothelial isoform of nitric oxide synthase (eNOS) by phosphorylation of serine 1177.^11, 12^ In humans and other mammals, despite high structural homology and activation of similar downstream pathways the biological processes regulated by insulin and IGF-1 are strikingly different.^13^ Consistent with this, in endothelial cells Kearney et al. demonstrated that deletion of IR reduced,^14, 15^ whereas deletion of IGF1R increased basal serine 1177 phosphorylated eNOS and insulin-mediated phosphorylation of serine 1177 on eNOS.^16^ They also showed that increasing IR in endothelial cells enhances insulin-mediated serine phosphorylation of Akt but blunts insulin-mediated serine 1177 phosphorylation of eNOS,^17^ whereas increased IGF1R reduces basal serine 1177 phosphorylated eNOS and insulin-mediated serine 1177 phosphorylation of eNOS.^18^

IR and IGF1R can bind insulin, IGF1 and IGF2, albeit with affinities such that they only bind their own cognate ligands at physiological concentrations.^19-21^ Due to the array of signalling pathways regulated by the IR and IGF1R, aberrant signalling through these receptors is associated with several diseases. Specifically, dysfunctional insulin signalling is the primary driver of type 2 diabetes mellitus, whilst altered IGF1 signalling manifests in several forms of cancer due to its role modulating cell proliferation.^22^

IR and IGF1R are unique amongst RTK’s in that they exist on the cell surface membrane as preformed dimers, contrasting with other RTK’s that only dimerise upon ligand binding. This means that the receptors share a unique mode of action amongst RTK’s; hormone binding elicits a large-scale conformational change to activate the receptor. The structural basis of activation of the insulin family receptors is still not fully understood, although recent findings have significantly contributed towards providing a plausible activation mechanism of the receptors.^23^ Ligands binding to the extracellular portion (ectodomain) of the receptors affect trans-autophosphorylation of the intracellular kinase domains, which in turn promotes binding and phosphorylation of adaptor proteins to affect subsequent downstream signalling.

Due to their high homology, the IR and IGF1R can heterodimerise to form functional hybrid receptors in tissues in which they are co-expressed, consisting of an IR monomer and IGF1R monomer. Hybrid receptors bind IGF1 with *ca.* fifty times higher affinity than insulin and *ca.* ten times higher affinity than IGF2.^19-21^ The physiological role, signalling properties and mechanisms regulating the formation of IR- IGF1R hybrid receptors are currently poorly understood. However, it is believed that hybrid receptors confer insulin resistance by sequestering IR protein and reducing the available insulin binding sites on the cell surface.^24, 25^ Similarly, increased numbers of hybrid receptors may increase the number of receptors binding IGF1 and IGF2, and therefore contribute to the signalling of these receptors in certain cancers.^26^ In skeletal muscle, fat and the heart, hybrid formation has been shown to exceed that of IGF1R and IR dimers.^27^ While the role of hybrids in human physiology is undefined there is a clear association with increased hybrids and situations of metabolic stress including: type 2 diabetes mellitus,^28, 29^ obesity,^30^ hyperinsulinemia,^31^ insulin resistance^32^ and hyperglycaemia.^33^Whilst there have been significant recent advances in the structural information available for the IR and IGF1R, limited structural information exists for hybrid receptors^34^, especially without IGF1 bound. Here we seek to further our understanding of hybrid formation using homology modelling and site-directed mutagenesis.

## Results and Discussion

### Homology Modelling of IR IGF1R Hybrid Receptors

As several related structures of the unliganded IR and IGF1R has been determined,^35, 36^ homology modelling provides a viable route for building a structural model of the apo-IR-IGF1R hybrid receptor. The published IR and IGF1R apo-receptor structures were examined to determine the major interfaces forming between the two receptor monomers. In both receptors, three separate interfaces form: between the first leucine-rich domain L1 and the second fibronectin type III domain FnIII-2’, the second leucine-rich domain L2: and the first fibronectin type III domain FnIII-1’ domains and between the α-chain of carboxyl terminal ⍺CT: first leucine-rich domain L1’ (where the apostrophe denotes a domain arising from the alternate monomer). With these interfaces present in both the IR and IGF1R structures, it was reasoned that the same interfaces were likely to form in the IR-IGF1R hybrid receptor. The TACOS^37^ (Template-based Assembly of COmplex Structures) server was initially chosen to produce the model as it is designed to model the structures of protein-protein complexes.

Sequence alignments were performed using MUSTER^38^ on residues 332-619 of the IR-B (UniProt ID: P06213-1) and 331-608 of the IGF1R (UniProt ID: P08069). TACOS identified the IR crystal structure (PDB:4ZXB) as the best template for the global protein complex, with 68% identity and 86% coverage. The normalised z-score for this alignment was 4.4, clearing the benchmark of 2.5 for a good alignment.

The interface between the IR and IGF1R is predicted to form between the β-sheet of the L2 domain and the second (C-terminal) β-sheet and loop regions of the FnIII-1 domains of each monomer respectively (Figure 1A, B). However, due to the differences in sequence of the IR and IGF1R chains in this region, the interactions are not truly conserved at the contacts between IR L2 and the IGF1R FnIII- 1, and vice versa. The predicted L2: FnIII-1’ interface is structurally similar to the analogous L2: FnIII-1’ interfaces of the IR^36^ and IGF1R^35^ unliganded ectodomain structures, with RMSD values of 1.149 Å and 0.832 Å relative to the relevant regions of the IGF1R (PDB: 5U8R) and IR (PDB: 4ZXB), respectively. Analysis of the TACOS model using PDBePISA^39, 40^ determined that the hybrid L2: FnIII-1’ interface covers an area of 1491.1 Å^2^, with a ΔG (Gibbs free energy) of −5.4 kcalmol^-1^. Typical PPIs exhibit interface areas ranging from 1300 to 2500 Å^2^ and free energies between −6 and −12 kcal mol^-1^ ^41^, meaning the hybrid L2: FnIII-1’ interface is both smaller and lower affinity than a typical PPI.

**Figure 1.**
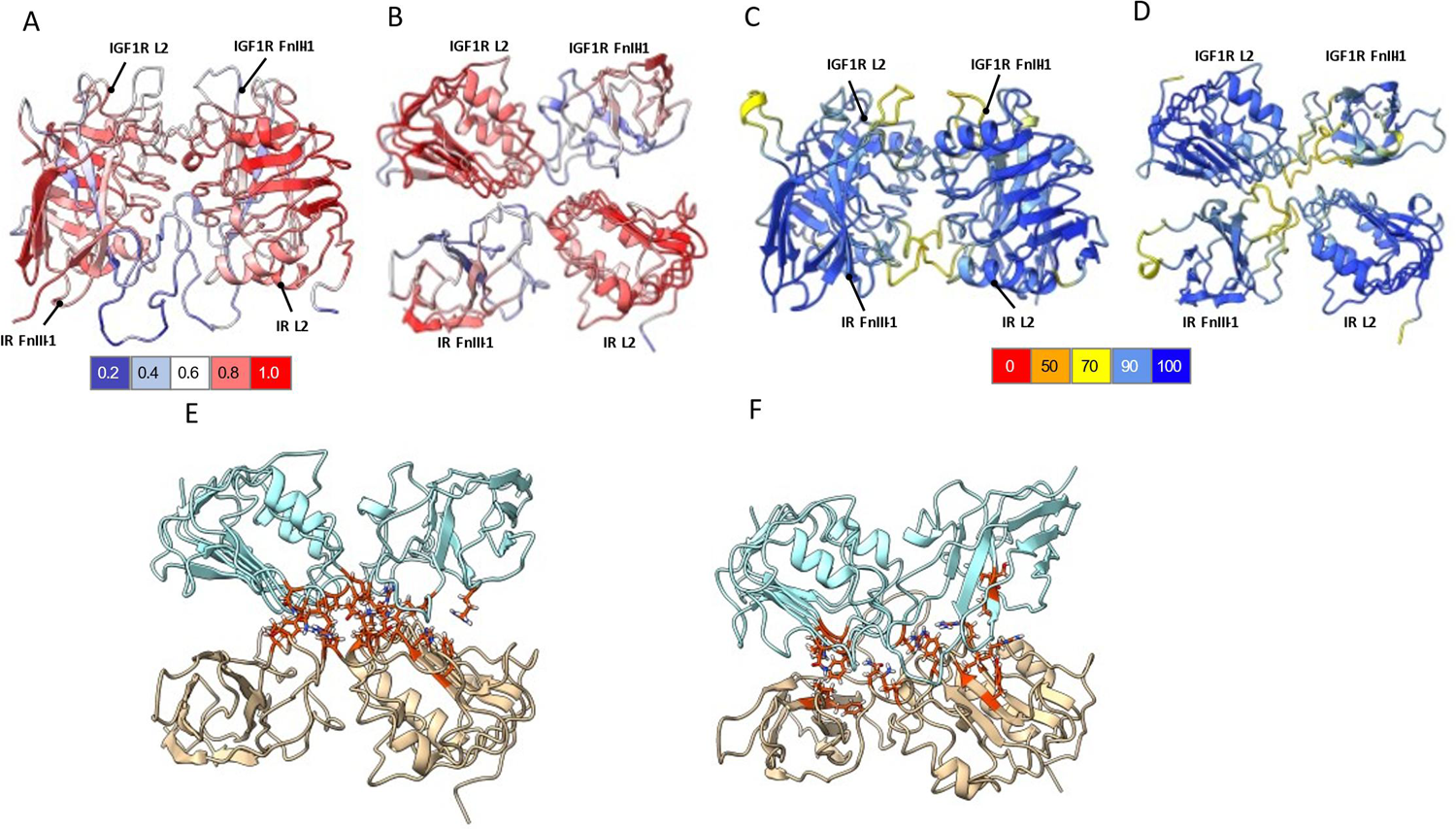
Homology models of the IR-IGF1R L2: FnIII-1’ interface. A) Face-on view of the TACOS homology model coloured by QMEANDisCO score according to the key (colour key at bottom); B) Top-down view of the TACOS homology model coloured by QMEANDisCO score (colour key at bottom). C) Face on view of the Alpha fold homology model with the ribbon coloured by pLDDT (colour key at bottom) D) Top-down view of the AlphaFold homology with ribbon coloured by pLDDT (colour key at bottom). E) KFC2 hotspot analysis of the Hybrid L2: FnIII-1’ interface for the TACOS homology model with IR shown as cyan ribbons and IGF1R as wheat-coloured ribbons. Hotspots identified in red. F) KFC2 hotspot analysis of the Hybrid L2: FnIII-1’ interface for the Alphafold homology model with IR shown as cyan ribbons and IGF1R as wheat-coloured ribbons. Hotspots identified in red.

More recently, the development of AlphaFold-multimer^42, 43^ for protein structure prediction provided an opportunity to further improve the homology model of the IR-IGF1R hybrid receptor. To allow direct comparison to the TACOS model, the sequences corresponding to residues 332-619 of the IR-B and 331-608 of the IGF1R sequences were modelled (Figure 1 C&D). The model predicted by AlphaFold showed good agreement to the TACOS model with an RMSD of 0.862 Å. Specifically, the secondary structure elements comprising the FnIII-1: L2 interface were highly concurrent, whilst the areas of lower agreement were typically located on the flexible loop regions of the FnIII-1 β-sheets. As the loop regions of the FnIII-1 β-sheets are lower resolution in the template crystal structures and likely highly flexible, discrepancies in modelling of these regions are not particularly concerning.

### Homology Model Validation

One limitation of the TACOS modelling methodology is the absence of integrated tools for assessing the quality of the generated models. Therefore, the SWISS-MODEL structure assessment suite^44^ was utilised to evaluate the TACOS model. QMEANDisCO^45^ score was used as the primary metric of assessment generating scores for both global and local quality evaluating the consistency of pairwise C⍺- C⍺ distances in the model by comparing them against restraints extracted from homologous structures allowing comparison of the model to an ensemble of experimental structures. The score is calibrated between 1 and 0, with below 0.6 indicating areas of lower quality.

The TACOS global QMEANDisCO score was calculated as 0.74 ± 0.05. Additionally, QMEANDisCO per residue analysis was utilised (Supplemental Figure 1) to assess the local quality relative to the overall model quality. Areas of significantly reduced local quality were identified at IR residues 533-554 and 581-594, as well as in IGF1R residues 544-566 and 599-614. These regions correspond to the β5 strand, β6 strand and connecting loop in the FnIII-1 domains. The lower quality of these regions is unsurprising, as these regions are relatively low resolution in both the template crystal structures. Additionally, a Ramachandran plot (Supplemental Figure 1) determined that 92.11% residues exhibited favoured dihedral angles, with 0.72% outliers. Overall, assessment metrics for the model indicated it was of good quality^44-46^, with specific regions in the FnIII-1 domain predicted to be of lower quality.

AlphaFold provides some additional metrics indicating the quality of regions of the predicted model. Firstly, AlphaFold calculates the predicted local distance difference test (pLDDT) which evaluates how well the environment in a reference structure is reproduced in a protein model through comparison of C⍺ local difference distance tests. The pLDDT for the AlphaFold model is generally high (Figure 1 C&D), with scores less than 70 isolated to the interdomain linker regions and flexible loops on the edges of the FnIII-1 domain. AlphaFold also produces a predicted aligned error (PAE)- a measure of the confidence in the relative positioning and orientation of domains in the AlphaFold prediction. The PAE for the AlphaFold model is generally lower than 10 Å (Supplemental Figure 2), indicating the confidence in the relative positions of the domains is high.

To allow direct comparison with the TACOS model, the AlphaFold model was also evaluated with the SWISS-MODEL structure assessment suite^44^. The AlphaFold model demonstrated a global QMEANDisCO score for the model was 0.78 ± 0.05, indicating a slight improvement compared to the TACOS model (Supplemental Figure 2). Consistent with the pLDDT score, per residue QMEANDisCO analysis identified lower quality regions of the FnIII-1 domains in both the IR and IGF1R domains. Importantly, these regions were confined to the same loop regions identified by the pLDDT score. This implies that the β5 and β6 strands comprising the core of FnIII-1 domain are predicted with significantly higher confidence in the AlphaFold model relative to the corresponding region of the TACOS model.

Ramachandran analysis of the AlphaFold model determined 96.29% residues occupied favoured conformations, with 0.53% outliers (Supplemental Figure 2). In comparison, the corresponding values for the TACOS model were 92.11% and 0.72% respectively. Overall, assessment of the AlphaFold and TACOS model indicates that the AlphaFold shows a modest improvement in structural quality, particularly in the β5 and β6 strands of the FnIII-1 domains.

### Prediction of Key Protein-Protein Interaction sites in Hybrids

Whilst PPI interfaces usually comprise a large area compared to a typical small-molecule binding pocket in a protein, it is accepted that a relatively small-number of amino acid residues contribute a large percentage of the total ΔG^f^ of a PPI interface^47^. Clusters of such residues are termed ‘hotspots’. The KFC2 server^48-50^ was chosen to computationally predict hotspot residues at the L2: FnIII-1’ interface of the TACOS model. KFC2 hotspot analysis of the Hybrid L2: FnIII-1’ interface classified a total of 27 residues as hotspots (Supplemental Table 1) which clustered at the top of the L2 domains of both chains (Figure 1E) with the interface formation largely governed by hydrophobic interactions, with no hydrogen bonds occurring between hotspot residues from either chain. In the mature hybrid receptor, the IR and IGF1R monomers are held together by intermolecular disulfide bonds, and the dominance of hydrophobic interactions at the L2:FnII-1 interface is consistent with that of many obligate PPI’s^51^. Specifically, residues IR Y457 and IGF1R F450 insert into hydrophobic cavities on the opposite monomer, forming critical reciprocal interactions. A further key cluster of contacts occurs where residues IR K487, T488, D491 and Q492 pack against IGF1R residues N478, R480, E484 and R485. The number and relative proximities of the predicted hotpot residues was encouraging in terms of disrupting hybrid formation, as they formed compact, well-defined regions.

Similarly, KFC2 hotspot prediction server was utilised to predict hotspot residues of the AlphaFold homology model. The KFC2a analysis predicted nine IR and nine IGF1R residues as hotspots respectively (Supplemental Table 2). These residues are clustered towards the top of L2 and FnIII-1 domains (Figure 1F), forming relatively localised epitopes.

The interactions mediated by the hotspot residues exhibit a degree of symmetry between the two receptor chains, although specific interactions are not always conserved. For instance, IR Y457 and IGF1R F450 establish reciprocal interactions by inserting into hydrophobic regions between the L2 and FnIII-1 domains. Similarly, both IR R398 and R399, as well as IGF1R R391 and H392 insert into charged grooves on the opposite FnIII-1 monomer to form electrostatic interactions. Additionally, both IR L196 and IGF1R I295 are involved in packing against hydrophobic regions on the opposite monomer.

Three hotspot residues from each chain were predicted by KFC2 for both the TACOS and AlphaFold model. These residues include IR Y457, IR K487, IR Q492, IGF1R F450, IGF1R N478 and IGF1R R485, which are all contained in the L2 domain. It is notable that there are discrepancies in the hotspot residues predicted within the FnIII-1 domains. This disparity can likely be attributed to reduced quality of this region in the TACOS model, indicating that the FnIII-1 residues predicted by the AlphaFold model should be prioritised over those from the TACOS model. The contrasting predictions of hotspot residues between the AlphaFold and TACOS models highlights the variations in local structural details despite the high RMSD indicating good overall agreement between the two models. This reinforces the importance of considering multiple modelling approaches and prioritising regions of higher confidence when utilising results obtained from homology models.

To investigate their potential functional significance, hotspot residues identified by KFC2 were compared with the Human Gene Mutation Database^52^, to determine if these residues had previously been identified as disease-causing mutations. Notably, the two point mutations IR K487E^53^, T488P^54^ have been documented as natural disease-causing variants associated with Donohue Syndrome. This disease is characterised by severe insulin resistance and significant growth restriction in humans, implying these residues have significant functional importance.

### Evaluation of Predicted Hotspots Using Site-Directed Mutagenesis

To experimentally validate the hotspot residues identified using the KFC2 server^48^, site directed mutagenesis was employed to generate a series of chimeric IR-IGF1R receptors containing point mutations at selected hotspot residues. Alanine mutations were chosen as they represent the truncation of the amino acid side chain to the β-carbon, whilst retaining the backbone dihedral angle preferences of most amino acids. These mutant receptors could then be characterised biochemically to determine the importance of each mutation to the IR-IGF1R PPI.

Mutants were generated by a PCR-based approach involving the amplification of the IR-Rluc and IGF1R-YPET plasmids (Supplemental Figure 3) utilising Platinum SuperFi II polymerase, which amplified IR-Rluc and IGF1R-YPET vectors specifically at a universal annealing temperature of 60 °C (Figure 2A). The IGF1R gene was sequenced to ensure the desired mutations had been incorporated and no additional mutations had been erroneously generated during PCR (Supplemental Figure 4).

**Figure 2.**
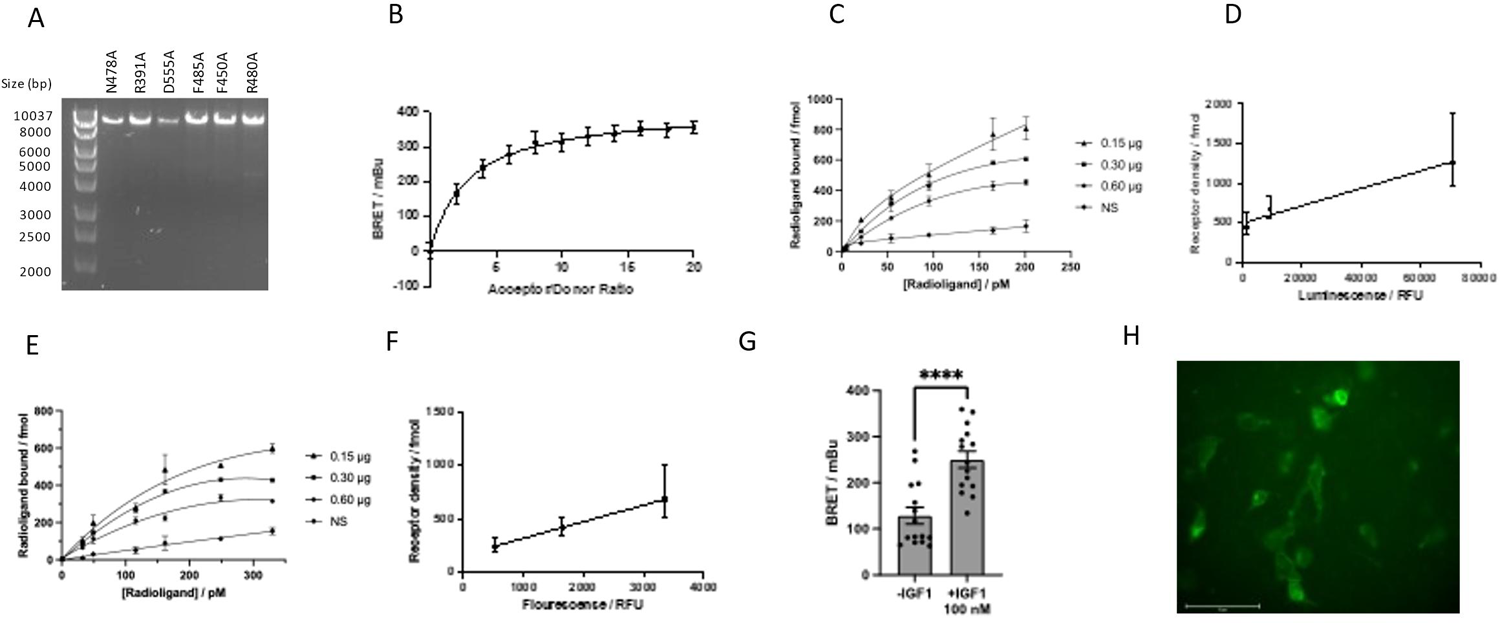
Radioligand binding assays to determine how IR-Rluc and IGF1R-YPET receptor quantities are correlated with their relative luminescence and fluorescence, respectively. A) PCR product when PCR was performed using SuperFi II Platinum mastermix. B) Donor saturation assay curve for BRET constructs IR-Rluc and IGF1R-YPET (±SEM). C) Saturation radioligand bind curves for the IR-Ruc constructs. The NS line depicts the non-specific binding component determined for untransfected cells (n=2; ± SEM. D) Calibration of IR-Rluc luminescence in the presence of 5 μM coelenterazine to receptor numbers (± SEM) E) Saturation radioligand bind curves for the IGF1R-YPET constructs. The NS line depicts the non-specific binding component determined for untransfected cells (n=2; ± SEM) F) Calibration of IGF1R fluorescence when excited at 513 nm to total receptor numbers (± SEM). G) BRET measurements showing the effect of treating cells co-transfected with IR-Rluc and IGF1R-YPET with IGF1 (100 nM, 5 min treatment, n=8,2; p<0.0001 ±SEM). H) Representative fluorescence microscopy images of HEK293 cells transfected with IR-Rluc and IGF1R-YPET, indicating the subcellar location of YPET, scale bar = 75 μm.

### Functional Evaluation of Hybrid Receptor Constructs

Once the mutated plasmid sequences had been verified, functional analysis of the resulting mutants was carried out. The ability of the mutant receptors to be transported in the cell membrane was evaluated by fluorescence microscopy, which detected the YPET labelled IGF1R hemi-receptor (Supplemental Figure 5). This experiment confirmed that yellow fluorescence was predominantly located at the cell membrane for wild-type IGF1R-YPET co-transfected into HEK293 cells with IR-Rluc. Similarly, each of the mutant receptor construct displayed fluorescence localised to the cell membrane when co-transfected with IR-Rluc. This contrasted to YPET transfected alone, in which high levels of fluorescence could be detected in the cell cytoplasm. Additionally, western blotting was utilised to ensure the mutant receptors signalled robustly in response to IGF1 stimulation (Supplemental Figure 6). Whilst this assay will also detect an IGF1R homomeric component, the presence of the mutant IGF1R constructs did not alter this signalling when compared to the signalling of the wild-type receptor. With confirmation that the mutated receptors did not show altered signalling or subcellular localisation, the effect of the mutations on the receptor binding interface could be evaluated.

To evaluate the effect of point mutations on hybrid formation, a cell-based bioluminescence resonance energy transfer assay based on the system previously described by Blanqaurt *et. al*^55^ was utilised (Supplemental Figure 7). The specificity of the BRET interaction between the IR-Rluc and IGF1R-YPET constructs was confirmed by a donor-saturation assay, in which the ratio of the BRET acceptor (IGF1R-YPET) was varied relative to the BRET donor (Figure 2B). The hyperbolic increase in BRET signal with increasing acceptor/donor ratio is typical of a specific BRET interaction, with the final plateau representing the saturation of all donor molecules.

To obtain receptor/donor ratios that equate to real protein quantities in the donor saturation assay, luciferase and YPET fluorescence must be correlated to receptor numbers to allow conversion. Receptor numbers for the IR-Rluc and IGF1R-YPET constructs were determined by a radioligand saturation binding assay. Cells were transfected with increasing quantities of cDNA encoding IR-Rluc or IGF1R-YPET. Subsequently, the luminescence and fluorescence values corresponding to IR-Rluc in the presence of 5 μM coelenterazine or YPET excited at 513 nm were determined for each transfection. Cells expressing the chimeric receptors were then probed with increasing concentrations of ^125^I-Insulin or ^125^I-IGF1 before bound probes were recorded. Non-specific binding was also evaluated by incubating cells with ^125^I-Insulin or ^125^I-IGFI in the presence of unlabelled insulin or IGFI at 500 nM to block specific binding (Figure 2 C&E). From these assays, the K_d_ for the IR-Rluc and IGF1R-YPET constructs were determined as 140 ± 25.2 pM and 122.8 ± 24.2 pM, which are in good accordance with the apparent K_d_ values of 190 pM and 120 pM reported^56^ for the IR and IGF1R respectively. It is noted in the case of IR-Rluc, the K_d_ value may be altered by the presence of the fused Rluc tag.

The B_max_ values for each transfection ratio were found to vary linearly with luminescence and fluorescence values for the IR-Rluc and IGF1R-YPET constructs respectively (Figure 2 D&F). This in turn allows the conversion between luminescence, fluorescence values and actual protein quantities for the IR-Rluc and IGF1R-YPET constructs respectively. It was reasonable to assume that point mutations in the receptor ectodomains would not affect luminescence and fluorescence originating from the C-terminal Rluc and YPET tags. Therefore, this analysis could similarly be utilised to convert between luminescence, fluorescence, and actual protein quantities for the chimeric mutant receptors.

Additionally, in cells transfected with IR-Rluc and IGF1R-YPET, an enhanced BRET signal could be observed upon stimulation with 100 nM of IGF1 (Figure 2G). This BRET increase was comparable in magnitude to that previously reported in hybrid receptors similarly tagged with Rluc and YPET^55^.

Finally, the subcellar location of the chimeric IR-IGF1R hybrid receptors was assessed to ensure they were correctly trafficked to the cell membrane. Using fluorescence microscopy, YPET signal was observed localised to the cell membrane in HEK293 cells co-transfected with IR-Rluc and IGF1R-YPET (Figure 2H). This result confirms that the attachment of BRET tags has not hindered the robust processing and trafficking of the chimeric hybrid receptors.

### Donor Saturation Assays to Characterise Hybrid Receptor Mutants

Donor saturation assays were undertaken by co-transfection of WT-IR-Rluc and mutant IGF1R-YPET, incorporating mutant IGF1R-YPET as part of a hybrid receptor. A non-linear regression model was used to fit to the data and calculate BRET_50_ and BRET_max_ values. The donor saturation assays determined that mutation of IGF1R residues F450A (Figure 3A), R391A (Figure 3B) and D555A (Figure 3C) did reduce the affinity of the IR IGF1R complex, as these mutants showed increased BRET_50_ values in the assay relative to the value for the hybrid WT receptor (Figure 3G). This result implies that each of these residues plays a significant role in stabilising the IR- IGF1R L2: FnIII-1’ interface.

**Figure 3.**
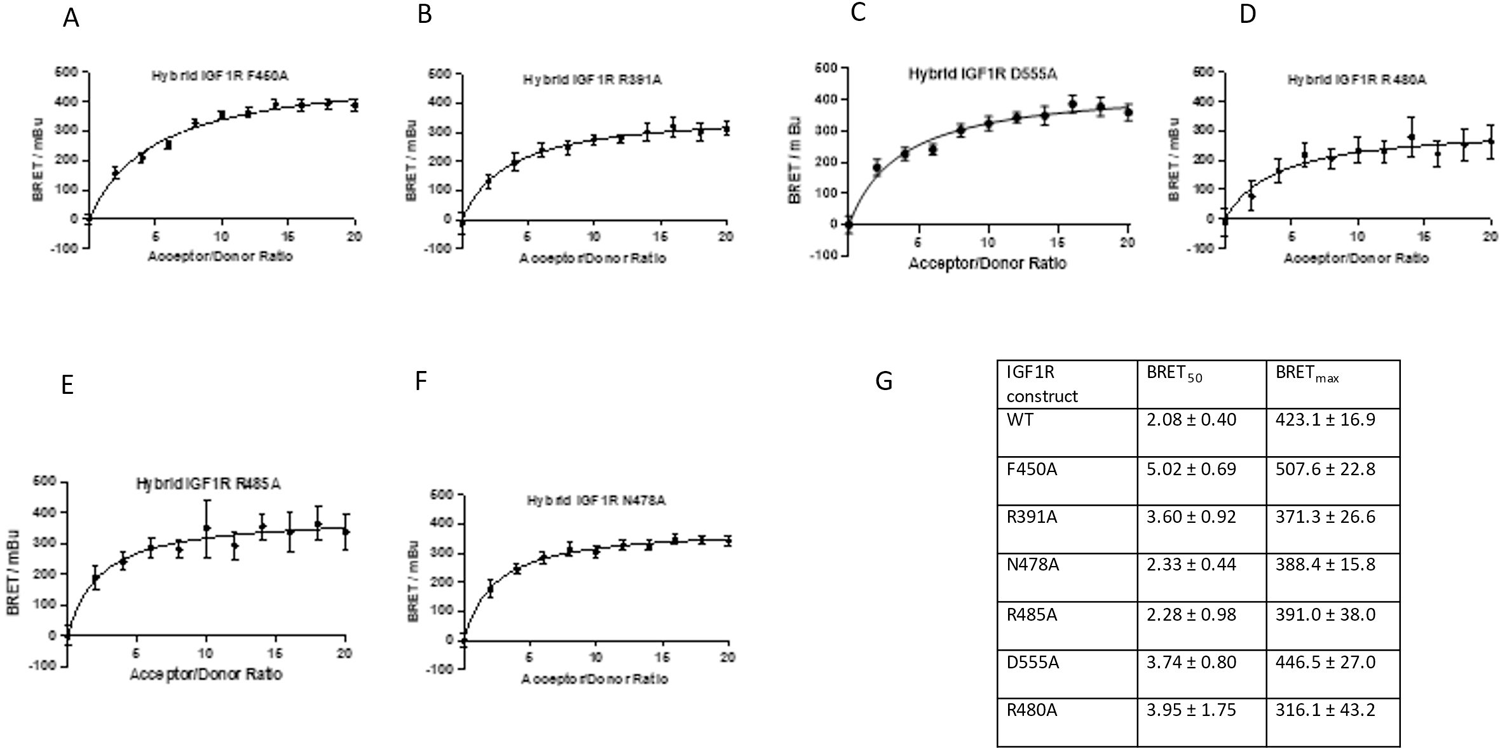
BRET_max_ and BRET_50_ parameters determined for each of the IR-IGF1R hybrid mutants in the donor saturation assay. A-F) Representative donor saturation assay curves performed for IR-IGF1R- hybrid mutants (n=8; ± SEM). G) BRET_50_ and BRET_max_ parameters determined for each of the IR-IGF1R hybrid mutants in the donor saturation assay. Error ranges represent the 95% confidence interval determined for the non-linear regression curve calculated from three separate experiments (n=8,3).

IGF1R residue F450 was predicted as a hotspot residue in both the AlphaFold and TACOS models (Figure 4A). In both models, the phenylalanine side chain inserts into a hydrophobic pocket on the IR monomer, stabilising the interface through hydrophobic interactions. Mutation of phenylalanine to alanine represents removal of the phenyl ring, which likely results in reduced hydrophobic contact and diminished interaction strength between the hybrid monomers. Noticeably, the mutant receptor also shows a substantial increase in BRET_max_ relative to the wild-type receptor, indicating that the mutation results in a change in the receptor conformation. As the BRET_50_ measurement is dependent to BRET_max_, the change in BRET_50_ cannot definitively be attributed to a change in IR and IGF1R affinity and may be due to the inherent relationship between the two parameters. Conversely, a change in receptor affinity could be affected by a large-scale conformational change of the receptor facilitated by the mutation F450A, which may explain the two-fold change in BRET_50_ from the point mutation of a single residue. It is notable that no change in subcellular location or signalling in response to IGF1 treatment was observed with this receptor. Therefore, any conformational change occurring due to the F450A mutation does not appear to substantially alter the receptors processing or signalling properties.

**Figure 4.**
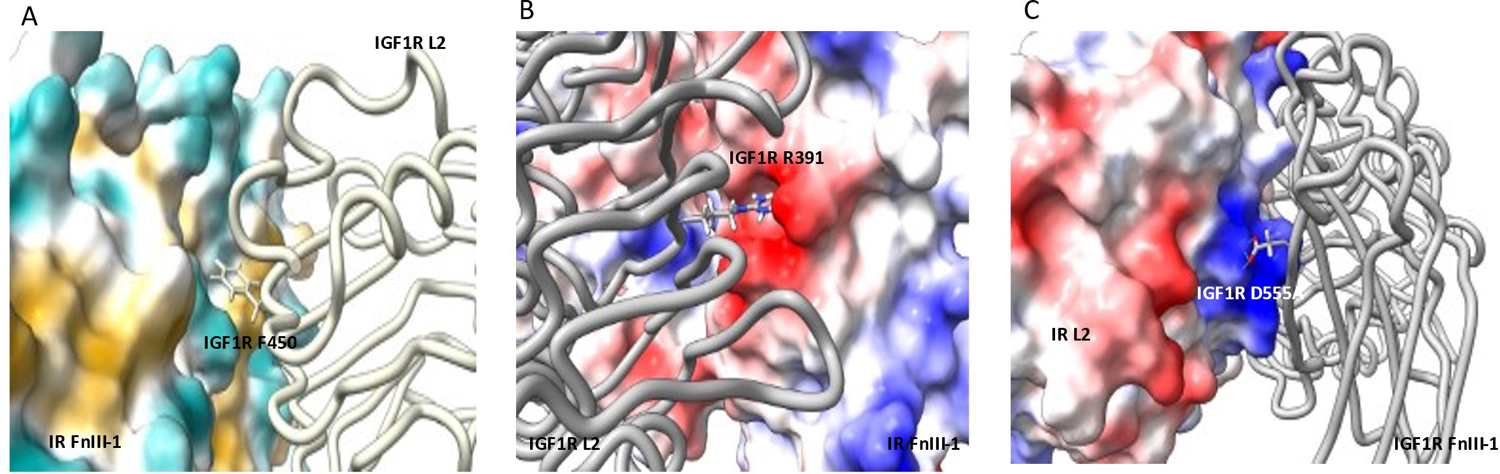
Predicted interactions of the IGF1R residues for which mutation to alanine cause an increase in BRET_50_. All images are taken from the AlphaFold homology model. A) Interaction of IGF1R F450 with the IR. The IGF1R is shown in yellow and IR with surface coloured by hydrophobicity; B) The interaction of IGF1R R391 with the IR. The IGF1R is shown in grey and IR with an electrostatic surface; C) The interaction of D555 with the IR. The IGF1R is shown in grey and IR with and electrostatic surface.

The residue R391 was predicted as a hotspot residue by the AlphaFold model and extends into a positively charged pocket on the IR monomer to form electrostatic interactions with surrounding residues (Figure 4B). Mutation to alanine results in the removal of electrostatic interactions and steric bulk and therefore reduce the affinity with the IR monomer. Similarly, residue D555 was predicted as a hotspot residue on by the AlphaFold model (Figure 4C). This residue extends to form electrostatic interactions with a cluster of residues positively charged residues located on the IR monomer.

Mutation to alanine would result in the removal of these electrostatic interactions. Both mutations show no significant change in BRET_max_ relative to the wild-type receptor, meaning that the change in BRET_50_ can be interpreted as a change in the receptor affinity upon mutation of the receptor.

The residues R480, R485 and N478 were similarly predicted as hotspot residues. However, there was no significant deviation in BRET_50_ observed with these mutants relative to the wild-type (Figure 3D-G). It is possible that these were hotspots, but the donor saturation assay was not sensitive enough to detect changes in affinity caused by these mutations.

The ability of residues IGF1R R391 and D555A to alter the apparent affinity of the hybrid receptor is significant, as it confirms that hybrid receptor formation can be modulated pre-dimerisation by changes to residues at the L2: FnIII-1’ interface. Once the receptors are dimerised, the formation of disulfide bonds effectively locks their dimeric arrangement^57^. Whilst mutations to residues at the L2: FnIII-1’ interface are unlikely to be the major cause of IR-IGF1R upregulation in response to metabolic disease^33^, the fact that receptor dimerisation can be modulated by this confirms epitopes at the L2: FnIII-1’ interface are a suitable target for therapeutic intervention.

## Conclusions

Two homology models of the IR-IG1F1R hybrid ectodomain were generated utilising TACOS and AlphaFold Multimer. These methodologies have independently predicted the formation of an interface between the L2 and FnIII-1 domains in the hybrid receptor. This interface was evaluated by SiteMap and KFC2, identifying suitable drug-binding regions and hotspot residues for small molecule intervention. Six IGF1R receptor mutants have been generated and their ability to disrupt hybrid formation has been assessed. All the generated mutants were expressed robustly, and the chimeric receptors IGF1R F450A, R391A and D555A all showed reduced affinity as part of a hybrid receptor.

There has been great progress in recent years towards the understanding of IR, IGF1R and hybrid structure and the conformational changes from apo to ligand bound receptors^36, 58-60^. In lieu of any current structural information regarding hybrid receptor apo structure, this work goes some way to understand the key hotspots for hybrid receptor dimerisation. This shows for the first time that hybrid receptor formation can be manipulated at the protein level and highlights the importance of the L2: FnIII-1’ interface in controlling hybrid receptor dimerisation. All the receptor mutants that showed reduced BRET_50_ relative to the wild-type receptor were predicted by the KFC2b method on the AlphaFold homology model, implying that this model can be used to predict hotspot features at the L2: FnIII-1’ interface with good accuracy.

A limitation of this approach is that the large size of the IR-IGF1R hybrid receptor precluded modelling of the entire receptor. Whilst it is likely that interactions between individual receptor domains is somewhat independent from the entire receptor, a complete model of the receptor would be preferable. It was also notable that the AlphaFold and TACOS models showed limited agreement on the identity of hotspot residues when evaluated by KFC2.

## Author Contributions

SJT, SPM and KJS designed and directed the study. SJT carried out the mutagenesis and analysed data. TI provided IR-Rluc and IGF1R-YPET plasmids for BRET assays. All authors, including MTK analysed the data and discussed the results. SJT and KJS prepared the manuscript.

