## Supplemental data for "Using site-directed mutagenesis to further the understanding of insulin receptor-insulin like growth factor-1 receptor heterodimer structure"

### Materials and Methods

#### *HEK293 Cell Culture*

HEK293 cells were stored under vapour phase liquid nitrogen. Cells were thawed by warming the cryovial in a water bath (37 °C) and seeding into prewarmed cell culture medium (10 mL) in a 75 $cm^2$ cell culture flask. A full medium change was undertaken once the cells had adhered, no longer than 24 hrs post thawing. Cells were passaged at 90% confluency. Cells were washed in prewarmed Dulbecco's Phosphate Buffered Saline (DPBS, Sigma Aldrich, 10 mL, 37 °C) before the addition of prewarmed trypsin (Gibco, 0.05%, 2 mL, 37 °C) and subsequently incubated for approximately 2 mins until detached from the flask. Prewarmed culture medium (37 °C) was added to achieve the appropriate seeding density for subsequent subculture. Cells were maintained in a 75 $cm^2$ cell culture flask in a humidified incubator (37 °C, 5% $CO_2$).

#### *Transfection of HEK293 cells*

HEK293 cells were seeded into a 6 well plate at 2.5 x $10^6$ cells per well and incubated (24 hrs) to achieve approximately 70% confluency by transfection. For transfections, Lipofectamine 2000 (Invitrogen) and cDNAs were used at a ratio of 5:1 v/w. To prepare the transfection mixture, Lipofectamine 2000 and the relevant cDNA were each diluted into OptiMEM (50 µL per well to be transfected) (Gibco). The diluted Lipofectamine solution was then mixed with the diluted cDNA and incubated (5 mins, rt). The resulting transfection mixture was then added dropwise to each well (100 µL) and the plate rocked to ensure even distribution. The transfected cells were then incubated for at least 12 hrs before harvesting or replating for further assays.

*BRET Assay*

HEK293 cells were seeded into 6-well plates at a density of 2.5 x $10^5$ cells per well and incubated. Cells were transfected as above. For experiments detecting hybrid receptors, cells were co-transfected with IR-Rluc cDNA and either IGF1R-YPET cDNA or empty pUC19 cloning vector cDNA to bring the total DNA to 0.6 μg per well. For experiments detecting homodimeric receptors, the combinations of IR-Rluc/ IR-YPET or IGF1R-Rluc/IGF1R-YPET were utilised, with the Rluc tagged construct co-transfected with empty pUC19. Transfected cells were incubated for 24 hrs. After 24 hours the transfected cells would typically be imaged using an Incucyte® Zoom (Sartorius) live cell imager to check the expression of YPET labelled protein and ensure transfection efficiency. Subsequently the cells were washed with DPBS (1 mL) (Sigma Aldrich) and the transfected cells pooled to reduce the effects of variation in well to well transfection efficiency. The cells were seeded into 96 well plate at a density of 1.5 x $10^4$ cells per well and incubated. After 24 hrs, cells were treated with relevant compounds in DMEM media (100 mM, 200 mL), diluted from DMSO stock (10 mM). After a further 24 hrs incubation, treated media was removed, wells were washed with DPBS (200 mL) and cells retained in DPBS (50 mL) for measurements. Plasmids encoding the constructs IR-Rluc, IGF1R-Rluc, IR-YPET and IGF1R-YPET were gifts from Dr. Tariq Issad (INSERM institute, Paris). Measurements were performed on a Envision® 2105 Multimode Plate Reader (Perkin Elmer). A solution of coelenterazine-h in ethanol (250 mM, 1 mL) was added to each well to give a final concentration of 5 mM in each well. Light-emission acquisition at 485 nm and 535 nm was started immediately. BRET signal was expressed in milliBRET unit (mBU). The BRET unit has been defined previously as the ratio 535 nm/485 nm obtained when the two partners are present, corrected by the ratio 535 nm/485 nm obtained under the same experimental conditions, when only the partner fused to *Renilla luciferase* is present in the assay. This can be represented as:

$$BRET = (E_{535} \div E_{485}) - Cf$$

Where $E_{535}$ corresponds to the luminescence at 535 nm, $E_{485}$ corresponds to the luminescence at 485 nm, and Cf corresponds to the ratio $E_{535}/E_{485}$ for the Rluc tagged construct transfected alone.

*IGF1 Treatment*

Cell culture and transfections were performed as above After 24 hrs post transfection, cells were changed into serum free DMEM (Gibco), containing no FBS supplement, and incubated (24 hrs). IGF1 (antibodies.com) dissolved in deionised water was added to each well at a final concentration of 100 nM and incubated (37 °C, 15 mins). Subsequently, the BRET ratio was measured as described above.

*Western Blotting Sample Preparation*

HEK293 cells were transfected with 0.3 µg of the relevant cDNA per well. After 24 hrs, cells were washed with ice-cold DPBS (1 mL) (Sigma Aldrich) and each well lysed by addition of lysis buffer (80 µL) (Invitrogen) on ice. The lysates were pre-cleared by centrifugation (14,000 g for 10 min at 4°C).

*Protein Quantification*

The protein concentration of cell lysates was determined using the Pierce BCA Protein Assay Kit (Thermo Scientific). Diluted albumin standards were prepared according to the manufacturers protocol. For each sample, 8 µL of protein lysate was diluted into 52 µL of water. 25 µL of diluted sample or albumin standard was plated into a clear 96 well plate in duplicate. The BCA working reagent was prepared by mixing BCA reagent A with BCA reagent B at a 50:1 ratio, and 200 µL of working reagent added to each sample or BCA standard. The plate was sealed and incubated for 30 mins at 37 °C. Subsequently, the absorbance of each well at 562 nm was determined. A standard concentration curve was generated from the albumin standards and cell lysate protein concentrations calculated by comparison to this concentration curve.

Gel Electrophoresis Procedures

*Immunoblotting*

50 µg of protein were mixed with NuPAGE™ LDS sample loading buffer (4X) (Invitrogen), NuPAGE™ sample reducing buffer (10X) (Invitrogen) and water to give the appropriate sample volume, before heating to 95 °C for 5 mins to ensure protein denaturation. Samples were loaded onto NuPAGE™ 4 to 12%, Bis-Tris gels (Invitrogen) alongside molecular weight markers samples and separated by SDS-PAGE electrophoresis for 90 mins at 110 V. Samples were then transfered to nitrocellulose membranes (Bio-Rad) using the TransBlot® Turbo Transfer System (Bio-Rad). The membrane was dried and incubated in 2% BSA TBST for 1hr to block non-specific binding. Membranes were treated overnight with primary antibodies at the relevant concentrations. Subsequently, the membranes were washed 3 x 10 mins in TBST before treatment with horseradish-peroxidase conjugated secondary antibodies of the relevant species (1:5000) in 5% BSA TBST for 1 hr. The membranes were then washed in 3 x 10 mins in TBST before visualisation.

*Protein Visualisation*

The immunoreactive proteins were visualised using the Western Chemiluminescent HRP substrate reagents (Immobilon) before imaging on the G:Box (Syngene) system. Blots were then stripped by application of stripping buffer for 15 mins before 3 x 10 mins of washes with TBST. Membranes were stored dry at 4°C. Images were analysed using the Image J software(Schneider, Rasband et al. 2012), with each band normalised to the signal of β-actin from the same sample for analysis.

*Fluorescence Microscopy*

HEK293 cells were maintained and transfected as above. Cells were transfected with IR-Rluc and empty pUC19 vector, or IR-Rluc and IGF1R-YPET mutants and incubated (24 hrs). The media was then replaced with OptiMEM (1 mL) (Gibco), and cells imaged on an EVOS cell imaging system (ThermoFisher) utilising the GFP filter.

*Mutagenesis*

Point mutations were integrated by including the desired nucleotide change into the forward primer, with at least 10 complimentary nucleotides present on the 3' end of the mutation. Reverse primers were designed so the 5' end of the forward and reverse primer anneal back-to-back. Primers were designed with NEBasechanger (https://nebasechanger.neb.com/) in the first instance. PCR was performed with Platinum™ SuperFi II PCR Master Mix (Invitrogen). PCR reactions were set up as follows:

| Reagent | Volume / µL | Final Concentration |
|---|---|---|
| 2X Platinum SuperFi II PCR mastermix | 25 | 1X |
| Nuclease Free Water | 19 | - |
| Forward Primer (10 µM) | 2.5 | 0.5 µM |
| Reverse Primer (10 µM) | 2.5 | 0.5 µM |
| Template DNA (1 ng$\mu L^{-1}$) | 1 | 1 ng total DNA |

Thermocycling conditions are detailed in below:

| Cycles | Step | Temp/ °C | Time / s |
|---|---|---|---|
| 1 | Initial denaturation | 98 | 30 |
| 25 | Denaturation | 98 | 5 |
| | Annealing | 60 | 10 |
| | Extension | 72 | 300 |

| | | | |
|---|---|---|---|
| 1 | Final extension | 72 | 300 |
| 1 | Hold | 4 | ∞ |

PCR reactions which resulted in a single band of the correct size were purified using QIAquick PCR purification kit (Qiagen). The purified PCR reaction (1 µL, 5-10 ng) was then added to a mixture of nuclease free water (5 µL) (NEB), 10X T4 Ligase buffer (1 µL) (NEB), T4 PNK (1 µL), T4 DNA ligase (1 µL) (NEB) and DpnI (1 µL) (NEB). The reaction was incubated at room temperature for 1 hr. 5α competent E-coli (50 µL) (NEB) stored at -80 °C were thawed on ice. Immediately upon thawing, reaction mixture (1 µL) was added to the 5α competent E-coli (50 µL). The sample was flicked 2-3 to mix and incubated on ice (30 mins). The sample was heat shocked (42 °C, 45 s) and then cooled on ice for a further 2 minutes. An aliquot of SOC media (950 mL) (NEB) was added to each sample and then incubate (37 °C, 1 hr, 220 rpm) on a benchtop incubator shaker. The final mixture (100 µL) was spread onto selective agar plates (100 µg $mL^{-1}$ AmpC) and incubated (37 °C, 16 hr). After incubation, single colonies were picked from the agar plates and used to inoculate overnight cultures in selective LB broth (10 mL, 100 µg $mL^{-1}$ AmpC) which were incubated (37°C, 16 hr, 230 rpm). An aliquot of the overnight culture (500 µL) was mixed with 50% glycerol in water (500 µL) and frozen at -80 °C to make a glycerol stock. The overnight cultures were centrifuged (4500 rpm, 4 °C, 10 min) and supernatant removed.

*Donor Saturation Assay*

HEK293 cells were seeded into a 12 well plate with 2 x $10^5$ cells per well. The cells were incubated for 24 hrs. The cells were transfected with plasmid using 110 ng of donor, increasing ratios of acceptor (x 0-20), and empty pUC19 maintain total DNA at 2130 ng per well (12 well). Transfections were performed at approximately 60% confluency. Plasmid DNA was diluted in OptiMEM (200 µL) (Gibco) and PEI (11.5 µL of a 1 mg $mL^{-1}$, 5:1 w/w PEI: DNA) (Polysciences) added to the DNA solution. The solution was immediately vortexed (5s) and incubated (RT, 15 mins). The transfection mixture (200 µL) was added dropwise to each well and the plate rocked to ensure even distribution over the well. The cells were incubated with the transfection reagents for 4 hrs, after which the media was exchanged for fresh media and the cells incubated for a further 44 hrs. Cells were then seeded at 5 x $10^5$ per well into white 96-well plate and incubated for 24 hrs. Simultaneously, the cells were seeded into a black 96 well plate. Cells were washed with DPBS (200 uL per well) (Sigma Aldrich) and resuspended in DPBS (50 µL). BRET measurements were recorded as above. Fluorescence measurements were recorded at 535 nm after excitation at 513 nm on cells in black plates.

*Radioligand Binding Assay*

HEK293 cells were seeded into 6-well plates at 2.5 x $10^6$ cells per well. After 24 hrs, the cells were transfected with IR-Rluc or IGF1R-YPET as above. Several transfections were performed with 0.15-0.6 µg cDNA per well. 24 hours later, cells were plated into a 96-well plate with 50,000 cells per well and incubated for a further 24 hours. Subsequently, the cells were washed with DPBS (100 µL per well) (Sigma Aldrich) and then maintained in DPBS (50 uL). For IR-Rluc transfected cells, coelenterazine-h was added to a final concentration of 5 µM and the luminescence recorded at 485 nm. For IGF1R-YPET transfected cells, the fluorescence at 535 nm was recorded for each well after excitation at 513 nm.

For radioligand binding, $^{125}$I-insulin or $^{125}$I-IGF1 (PerkinElmer) were diluted to the working concentration with unlabelled insulin or IGF1 respectively. This was then further diluted in DPBS to give six separate ligand concentrations spanning 0.5-10 times the Kd value for the relevant receptor-ligand complex. An aliquot of each radioligand was then removed and unlabelled insulin or IGF1 added at a final concentration of 500 nM (*ca.* 100 x Kd) to allow the measurement of non-specific radioligand binding. A further aliquot of radioligand was retained to determine the specific activity of each radioligand concentration. The radioligand was then added to the cells in duplicate wells for each concentration value and the cells incubated (3 hrs, 37 °C). Post-incubation, the cells were washed with ice-cold DPBS (100 µL) and lysed by addition of cell lysis buffer (50 µL) (Invitrogen). Cell lysates were removed into counting vials, and each well washed further lysis buffer (50 µL). Vials were counted on an AMG Automated Gamma Counter (Hidex). The $K_d$ and $B_{max}$ values were determined using GraphPad Prism using non-linear regression model with single site fitting.

*TACOS Model Generation*

Sequences for residues 332-619 of the human IR-B and 331- 608 of the human IGF1R downloaded from the UniProt Databse(Bienert, Waterhouse et al. 2016, UniProt 2021, Consortium 2022). These were submitted to the TACOS online structure prediction server (https://zhanglab.ccmb.med.umich.edu/TACOS/) with default settings.

*AlphaFold Model Generation*

Sequences for residues 332-619 of the human IR-B and 331- 608 of the human IGF1R downloaded from the UniProt Databse(Consortium , UniProt 2021, Consortium 2022). These were submitted to the ColabFold(Mirdita, Schütze et al. 2022) using Alphafold2 multimer(Evans, O'Neill et al. 2021) using 20 recycles with tolerance = 0.05. pLDDT values and the PAE plot were downloaded from the results file UCSF ChimeraX(Pettersen, Goddard et al. 2021) used to visualise the resulting model.

*Homology Model Quality Assessment*

Models were submitted to the SWISS-MODEL (https://swissmodel.expasy.org/) structure assessment server(Studer, Rempfer et al. 2020). QMEANDisCO values were downloaded and visualised in UCSF ChimeraX(Pettersen, Goddard et al. 2021).

*Hotspot Prediction*

Hotspot prediction using the resulting IR-IGF1R hybrid models was performed using KFC2 server (https://mitchell-web.ornl.gov/KFC_Server/index.php)(Zhu and Mitchell 2011). Visualisation of the homology model and hotspot analysis was performed using UCSF ChimeraX(Pettersen, Goddard et al. 2021).

*Statistics*

Data were analysed using GraphPad Prism software (version 9)[195]. P values > 0.05 were deemed statistically significant. Results are expressed as mean ±SEM unless otherwise stated.

Comparisons between two groups were performed using an unpaired Student's t-test. Comparisons between the mean values of multiple groups were performed using one-way analysis of variance (ANOVA), followed by Tukey's multiple comparisons test. Non-linear regression curves were calculated using a single-site fitting model, accounting for non-specific binding if appropriate.

<u>Supplemental Figures</u>

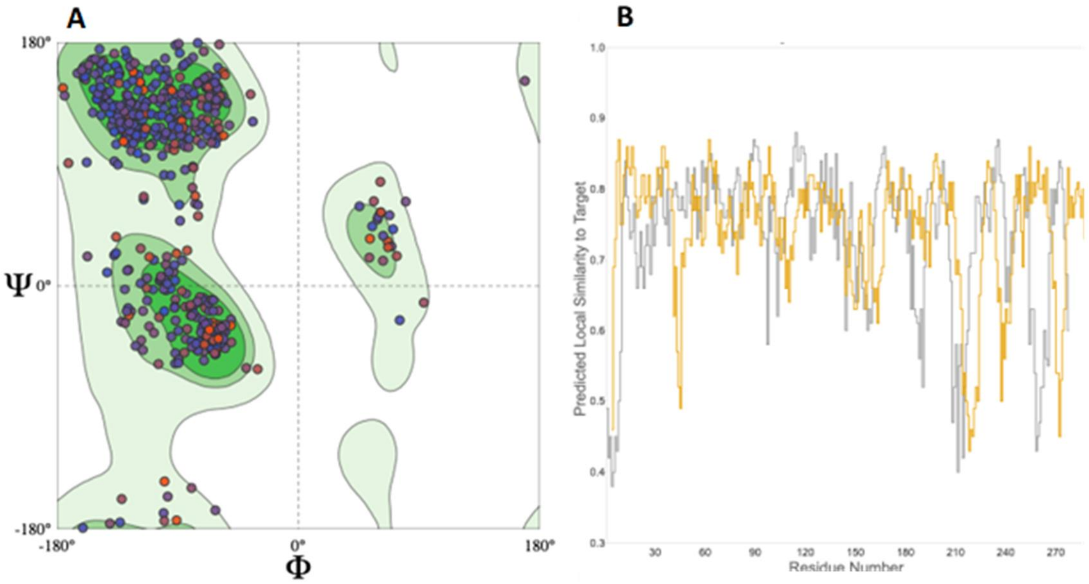


Figure S1-TACOS homology model validation with SWISS-MODEL structure assessment; A) Ramachandran plot of TACOS homology model. ϕ = C-N-Cα-C angle, Ψ = N-Cα-C-N angle of amino acid backbones. Contours of favoured regions are extracted from 12,521 non-redundant experiment structures (pairwise identity cut-off 30%, resolution cut-off 2.5 Å) such that 99.7% ϕ/Ψ are contained in the first contour, 95.0% in the second contour and 80.0% in the third; B) Plot of the per residue QMEANDisCO score. The IR monomer (grey) and IGF1R monomer (orange) are both shown. Residue numbers refer to model numbering.

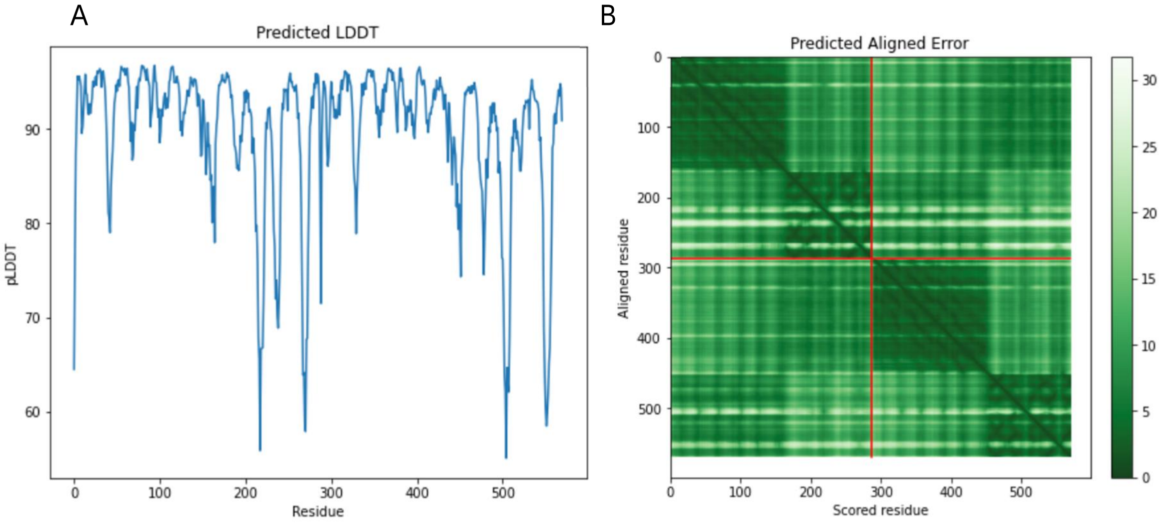


Figure S2- AlphaFold homology model of the L2: FnIII-1 interface. A) Plot of the per residue pLDDT score; B) Plot of the PAE for the homology model.

| Chain | Amino acid | Residue | KFC2a | KFC2a Conf | KFC2b | KFC2b conf |
|---|---|---|---|---|---|---|
| IR | GLN | 433 | Hotspot | 0.48 | ------- | -0.07 |
| IR | PHE | 454 | ------- | -0.03 | Hotspot | 0.02 |
| IR | HIS | 456 | Hotspot | 0.71 | Hotspot | 0.10 |
| IR | TYR | 457 | Hotspot | 0.47 | ------- | -0.31 |
| IR | PRO | 459 | Hotspot | 0.92 | ------- | -0.37 |
| IR | LYS | 487 | Hotspot | 1.59 | ------- | -0.02 |
| IR | THR | 488 | Hotspot | 1.18 | ------- | -0.29 |
| IR | ASP | 491 | Hotspot | 1.88 | ------- | -0.12 |
| IR | GLN | 492 | Hotspot | 1.58 | Hotspot | 0.05 |
| IR | THR | 598 | Hotspot | 0.99 | ------- | -0.50 |
| IR | SER | 600 | Hotspot | 0.37 | ------- | -0.61 |
| IGF1R | PHE | 450 | Hotspot | 0.62 | Hotspot | 0.04 |
| IGF1R | PRO | 452 | Hotspot | 0.77 | ------- | -0.42 |
| IGF1R | LYS | 453 | Hotspot | 0.04 | ------- | -0.33 |
| IGF1R | ASN | 478 | Hotspot | 0.23 | ------- | -0.42 |
| IGF1R | ARG | 480 | Hotspot | 1.13 | Hotspot | 0.04 |
| IGF1R | ASN | 481 | Hotspot | 1.15 | ------- | -0.06 |
| IGF1R | GLU | 484 | Hotspot | 0.79 | ------- | -0.26 |
| IGF1R | ARG | 485 | Hotspot | 1.40 | Hotspot | 0.28 |
| IGF1R | ARG | 518 | Hotspot | 0.42 | ------- | -0.18 |
| IGF1R | LEU | 586 | Hotspot | 1.09 | ------- | -0.03 |

Table S1 Hotspot residues predicated by KFC2 for the TACOS homology model. Residues also predicted by KFC2a in the AlphaFold homology model are highlighted in blue.

| Chain | Amino acid | Residue | KFC2a | KFC2a Conf | KFC2b | KFC2b Conf |
|---|---|---|---|---|---|---|
| IR | ARG | 398 | Hotspot | 0.47 | Hotspot | 0.23 |
| IR | ARG | 399 | Hotspot | 1.11 | Hotspot | 0.15 |
| IR | TYR | 428 | Hotspot | 0.17 | Hotspot | 0.14 |
| IR | LEU | 430 | Hotspot | 1.43 | Hotspot | 0.29 |
| IR | TYR | 457 | Hotspot | 0.99 | Hotspot | 0.11 |
| IR | LYS | 487 | Hotspot | 0.53 | ------- | -0.52 |
| IR | GLN | 492 | Hotspot | 0.33 | ------- | -0.35 |
| IR | LEU | 528 | Hotspot | 0.57 | ------- | -0.13 |
| IR | PHE | 530 | -------- | -0.52 | Hotspot | 0.04 |
| IR | LEU | 596 | Hotspot | 0.85 | ------- | -0.02 |
| IGF1R | ARG | 391 | Hotspot | 0.46 | Hotspot | 0.13 |
| IGF1R | HIS | 392 | Hotspot | 0.85 | Hotspot | 0.17 |
| IGF1R | LEU | 423 | Hotspot | 0.93 | Hotspot | 0.24 |
| IGF1R | PHE | 450 | Hotspot | 1.16 | Hotspot | 0.23 |
| IGF1R | ASN | 478 | Hotspot | 0.13 | ------- | -0.5 |
| IGF1R | ARG | 485 | Hotspot | 0.43 | ------- | -0.18 |
| IGF1R | ILE | 521 | Hotspot | 0.1 | ------- | -0.29 |
| IGF1R | SER | 522 | Hotspot | 0.42 | ------- | -0.42 |
| IGF1R | ASP | 555 | Hotspot | 0.62 | Hotspot | 0.05 |

Table S2 Hotspot residues predicted by KFC2 for the AlphaFold homology model. Residues also predicted by KFC2a in the TACOS homology model are highlighted in blue.

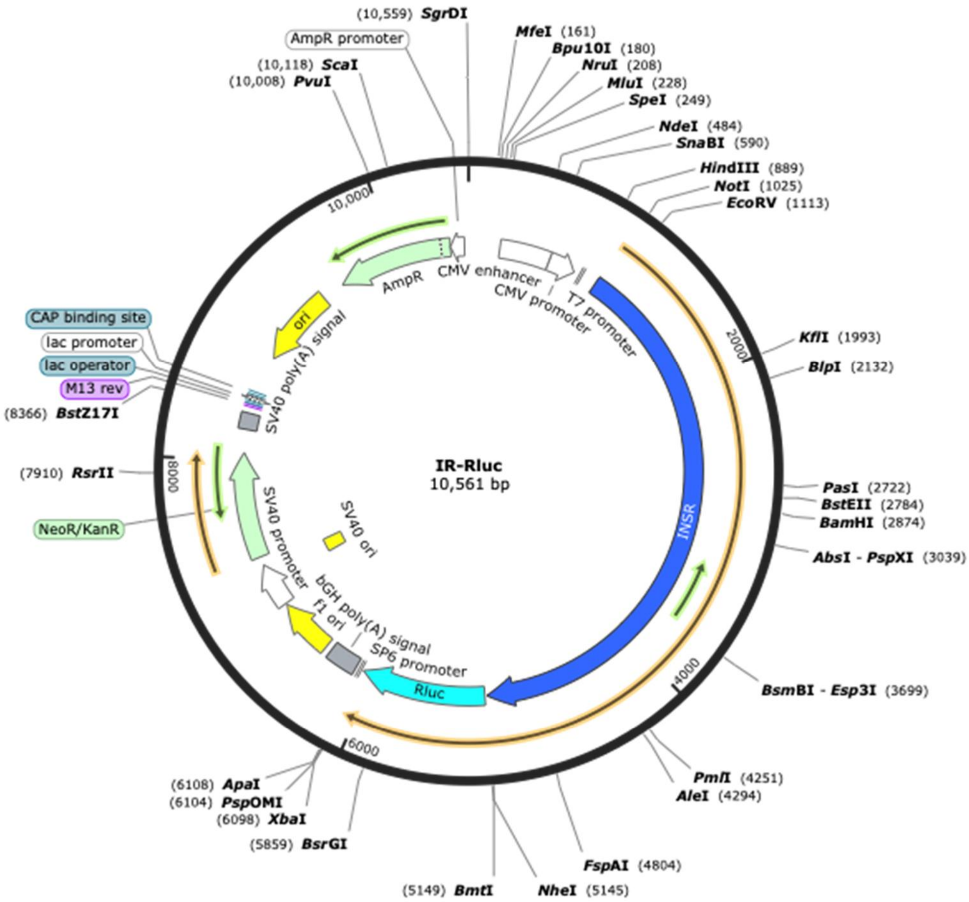

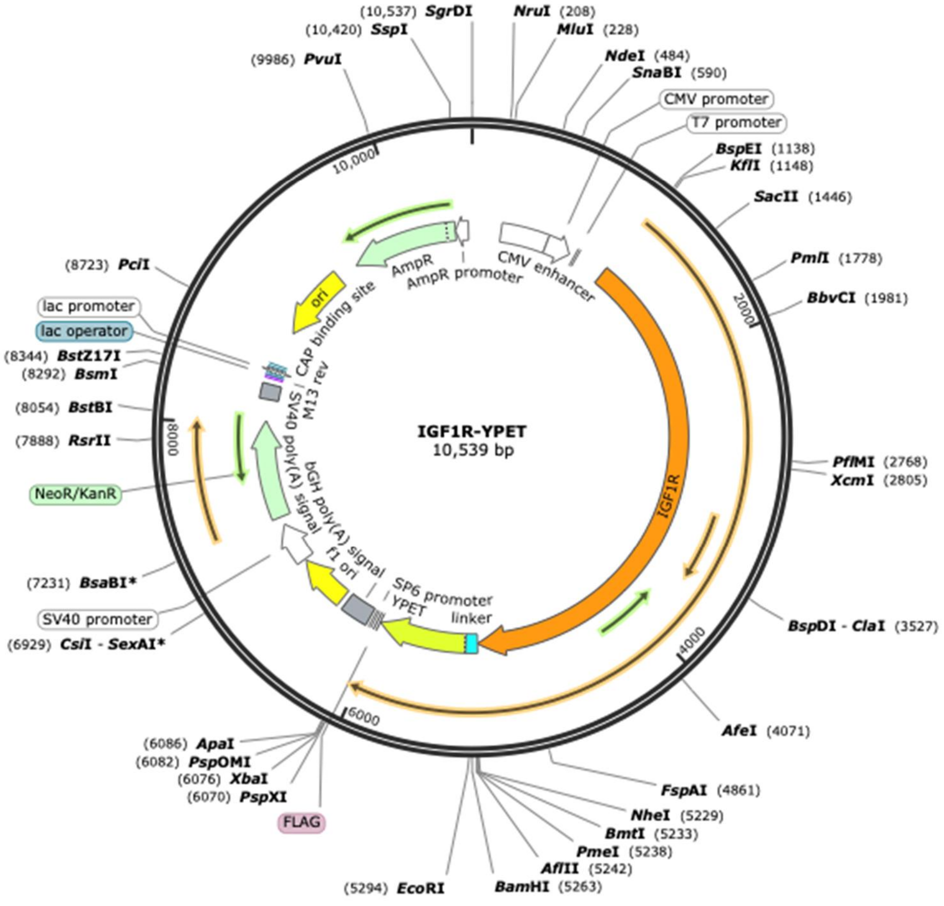


Figure S3 Plasmid maps for IR-Rluc (top) and IGF1R-YPET (bottom)

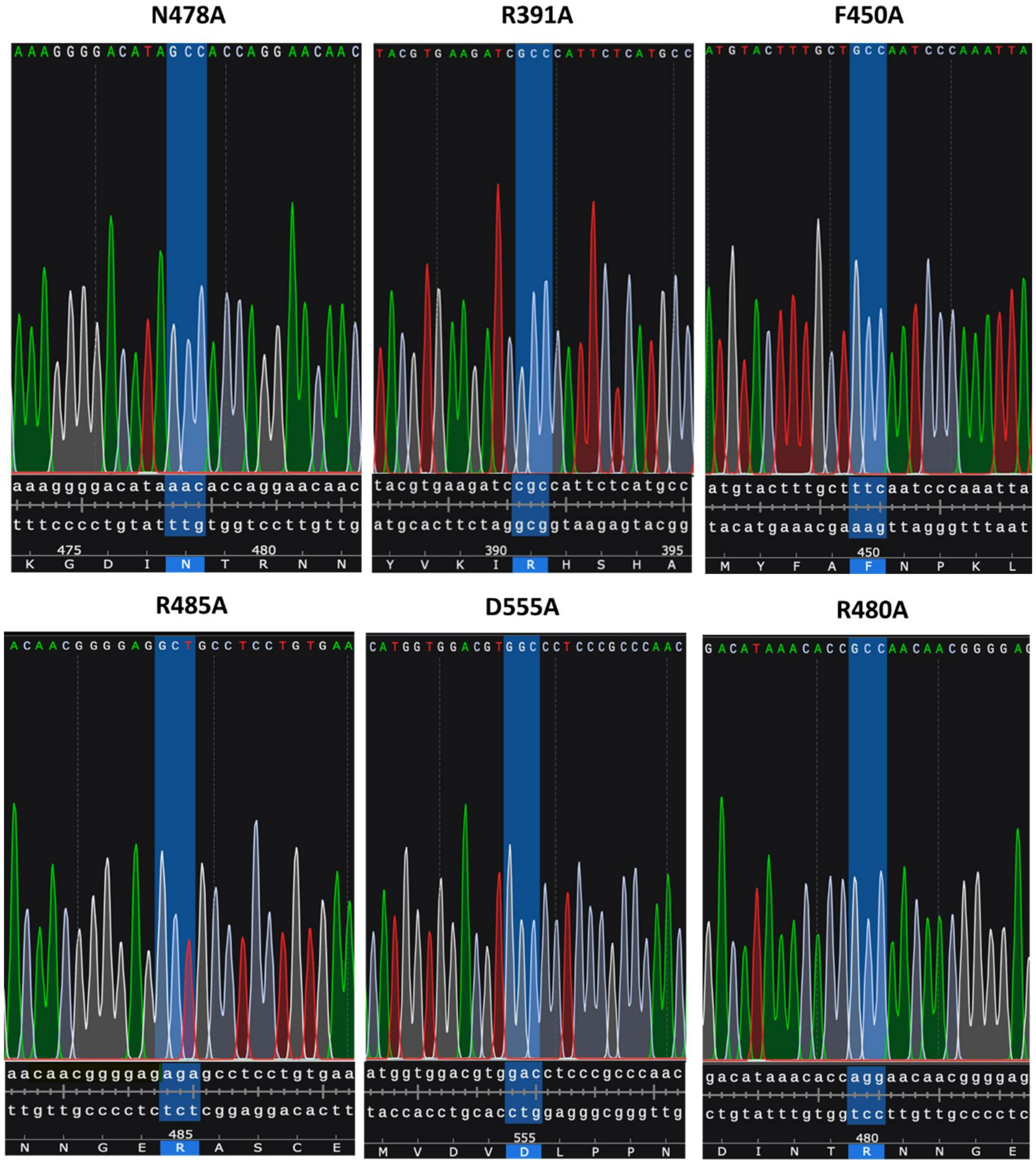


Figure S4 Sequencing of mutant vectors to verify point mutagenesis. Mutant sequencing traces are shown above, and relevant region of the wild-type sequences are shown below.

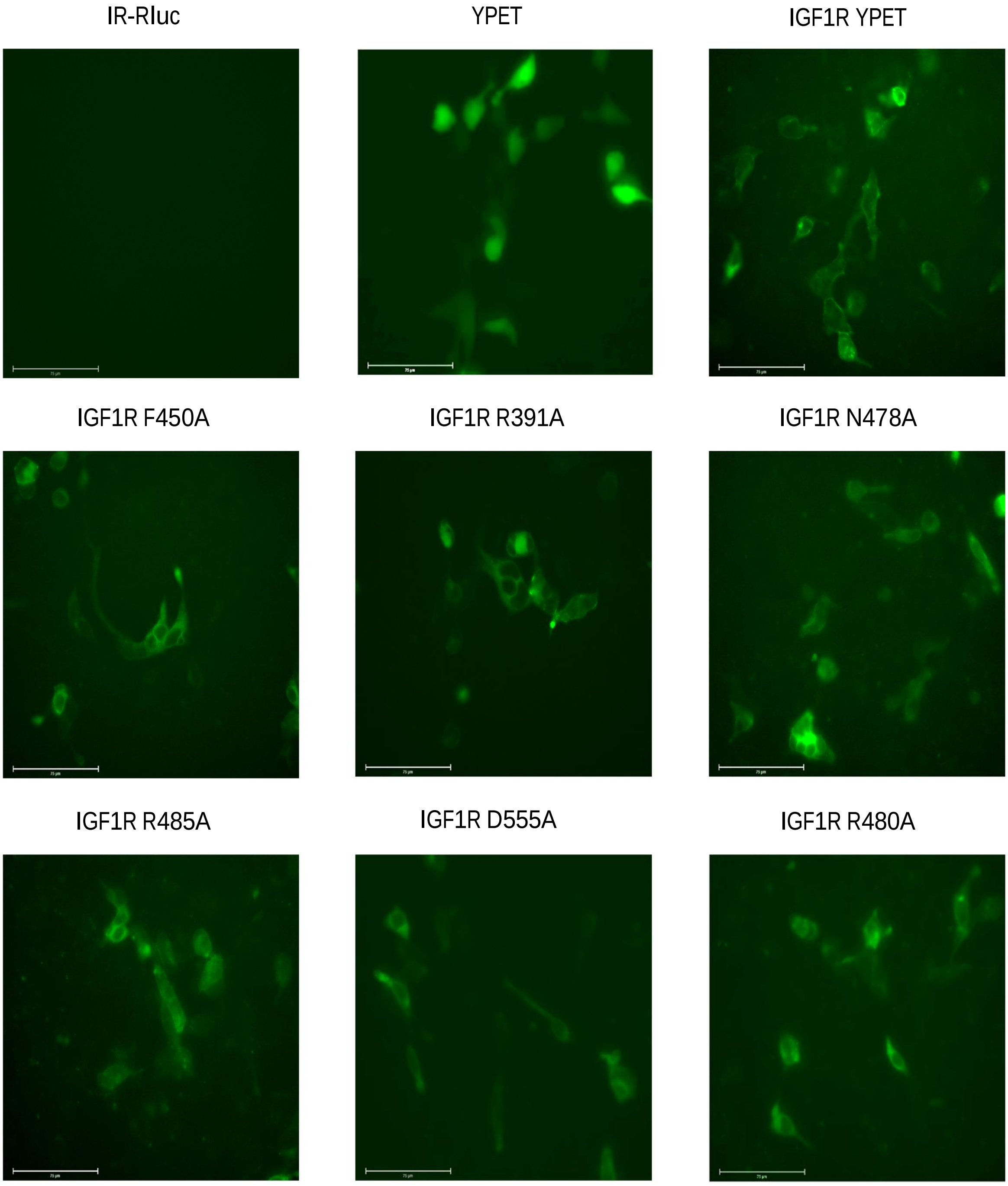


Figure S5 Representative fluorescence microscopy images of cells transfected with combinations of IR-Rluc and the indicated IGF1R mutants construct, showing the subcellular location of the YPET fusion protein. Scale bar = 75 µm.

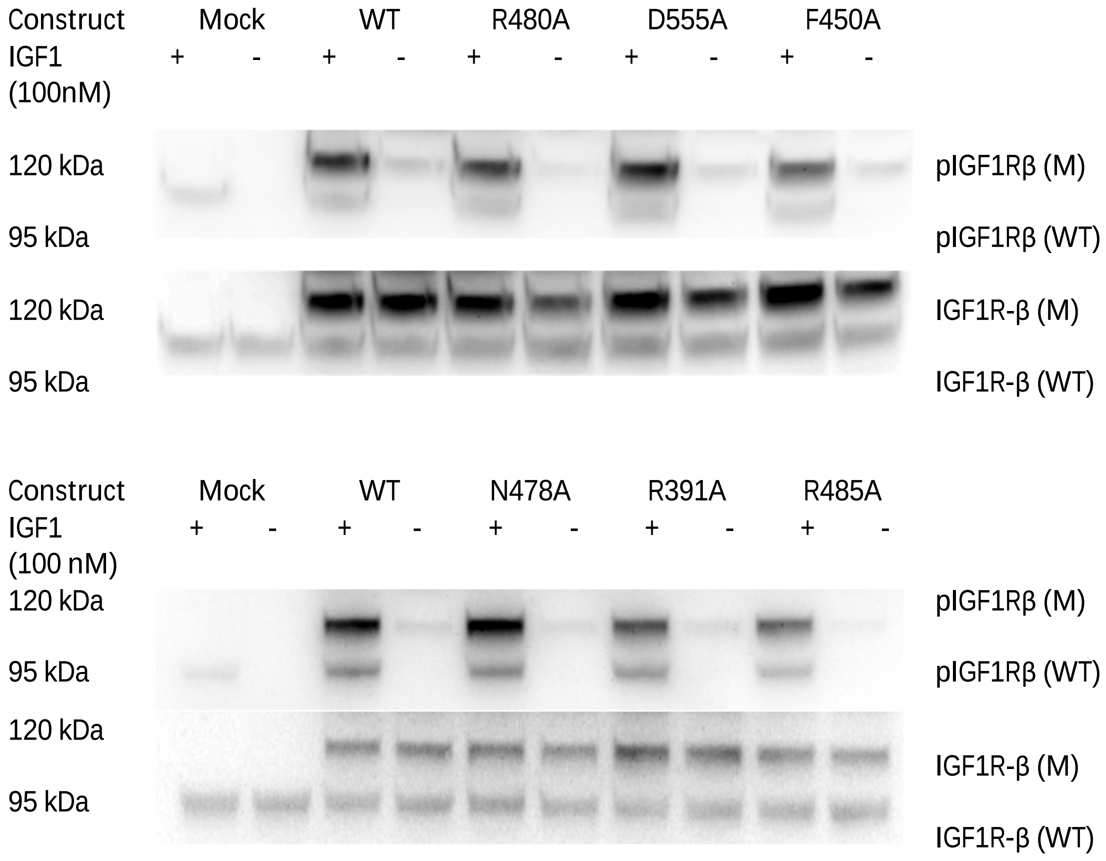


Figure S6 Representative western blots of HEK293 cells transfected with combinations of IGF1R-YPET mutants and treated with 100 nM IGF1, showing the levels of IGF1R pY1135 and total IGF1R when probed by the pIGF1R specific DA7A8 antibody (top) and the IGF1R specific D23H3 antibody (bottom). The labels (M) and (WT) refer to the mutant and wild-type bands respectively.

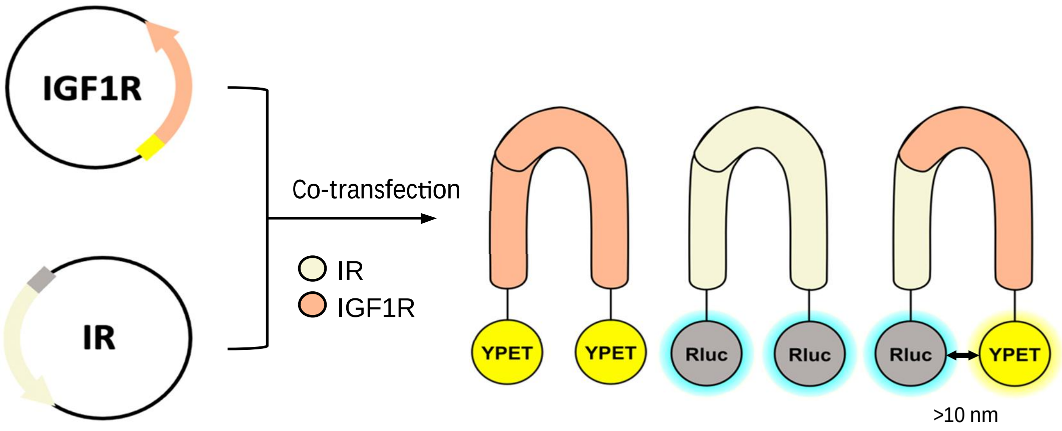


Figure S7 Schematic of BRET assay to specifically detect IR-IGF1R hybrid formation in live cells. Plasmids encoding the IR and IGF1R tagged with Rluc and YPET respectively are transfected into HEK293 cells, allowing detection of hybrid receptors by a RET interaction between the Rluc and YPET tagged monomers.

Raw Image Files

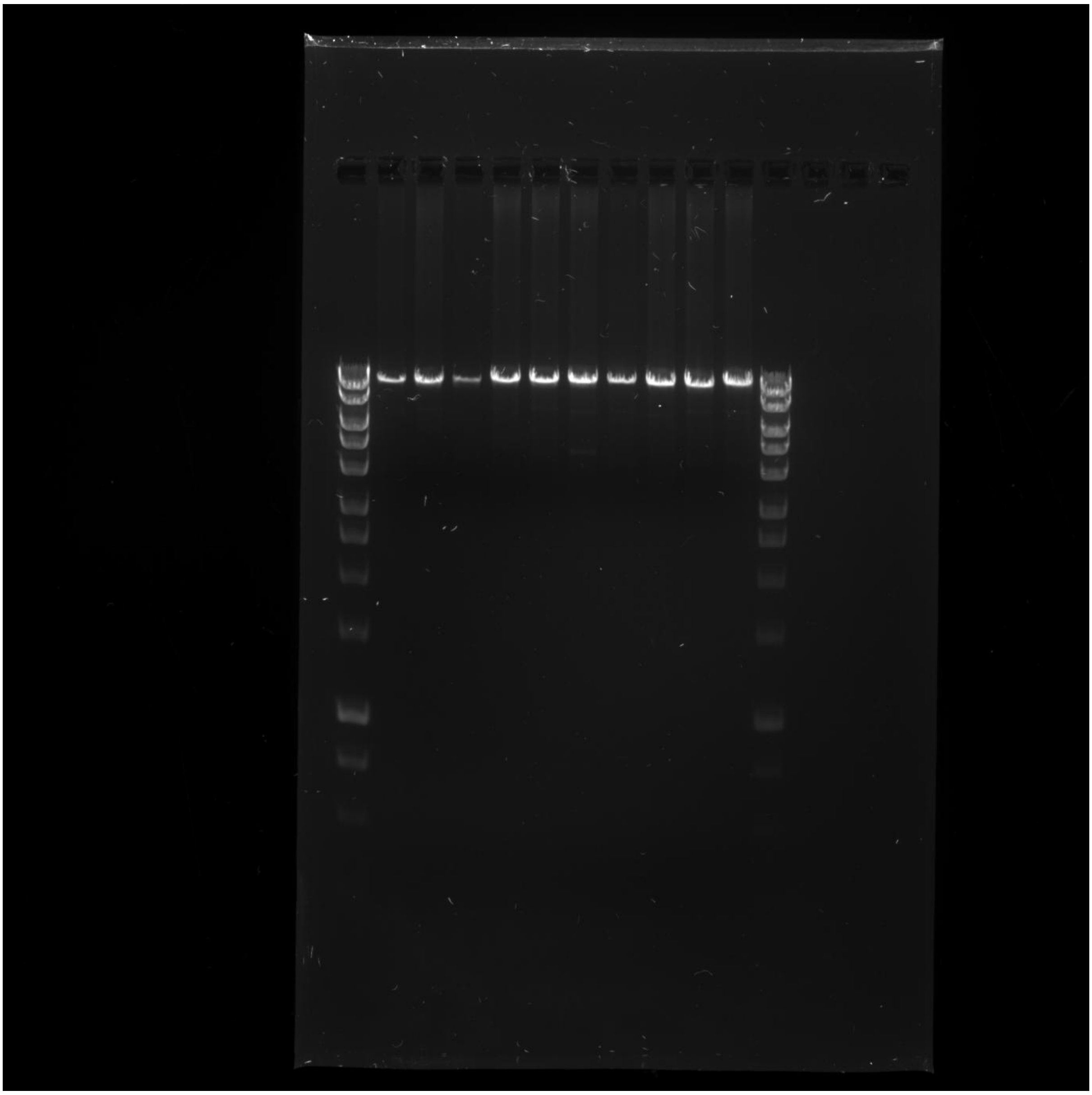

Raw image file of Figure 2A

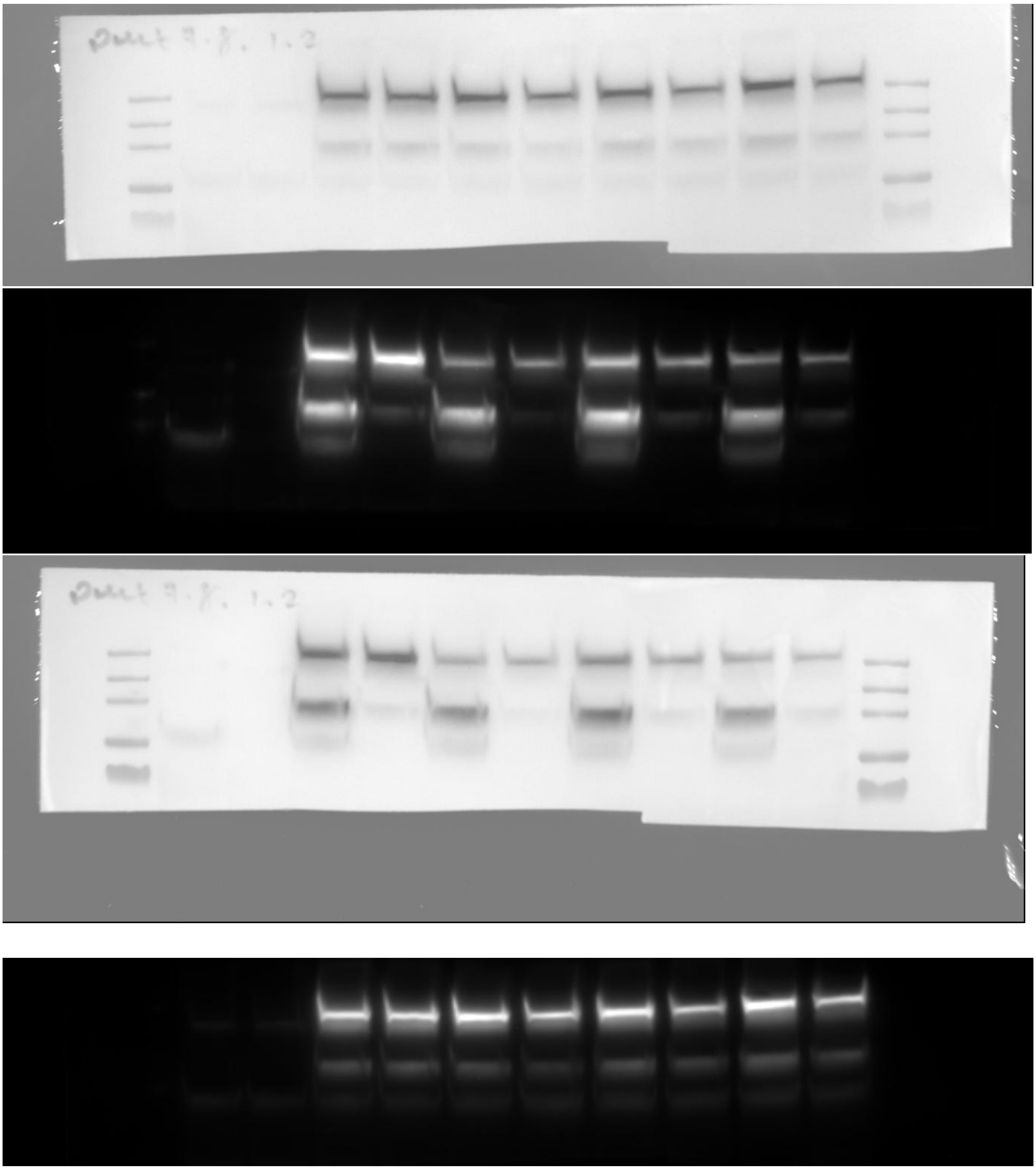

Raw image files of Figure S6
